# Dynamic Patterns of Nuclear Transcription Factor Abundance in Plant Basal Immunity Revealed by Spatial Proteomics of Arabidopsis Nuclei

**DOI:** 10.64898/2026.06.30.735533

**Authors:** Mohamed Ayash, Carsten Proksch, Domenika Thieme, Nicole Bauer, Justin Lee, Ingo Heilmann, Wolfgang Hoehenwarter

**Author notes:** Author for correspondence: Wolfgang Hoehenwarter Alternative.

## Abstract

- The control of amount of nuclear proteins is fundamental in regulating plant gene expression, but the mechanisms of quantitative dynamics of the nuclear proteome are largely unstudied during adaptive responses to pathogens.
- Highly specific labeling, enrichment and measurement of the nuclear proteome was performed using TurboID LC-MS of *Arabidopsis thaliana* leaves treated with the pathogen-associated molecular pattern (PAMP), flg22, and/or cycloheximide. The chosen experimental approach allowed discrimination of the effects of translation, nuclear protein import, trafficking of preexisting proteins, derepression, and nuclear protein turn-over upon elicitation of basal immunity.
- The highly specific, deep coverage of proteins in the nucleus makes this study a resource for anyone interested in plant nuclear proteome dynamics and defense.
- Around 2,000 nuclear proteins were repeatedly quantified, including more than 300 transcription factors or other proteins related to transcription. Several proteins with documented activity in endosomes were newly synthesized and imported into nuclei upon PAMP challenge, suggesting alternative nuclear functions in PAMP-triggered immunity (PTI). Circadian clock components, including the transcription factor, CIRCADIAN CLOCK ASSOCIATED 1 (CCA1)-HIKING EXPEDITION (CHE), were depleted upon PAMP challenge, suggesting a safeguard against untimely induction of systemic acquired resistance (SAR).
- Based on proteomic patterns, proteins moonlighting in the nucleus as well as trafficking and turn-over regulation of the proteome are common elements during plant immunity.

## Introduction

Gene expression patterns evolve rapidly as plants run genetic programs in development and interact with a continuously changing environment. To a large part gene expression patterns are shaped by transcriptional and co-/post-transcriptional processes in the cell nucleus. Transcription factor (TF) and co-regulator activity, 3-dimensional chromatin state and its remodeling, mRNA modification and alternative transcription initiation and/or splicing are essential elements contributing to the apparent patterns of gene expression. The nuclear proteins involved in all of these processes are targeted by intra-cellular signaling that communicates the need for physiological states that represent appropriate responses to a perceived stimulus.

Plant immunity (Jones *et al*., 2024) can be elicited at the cell surface by the perception of pathogen-associated molecular patterns (PAMPs) either by pattern recognition receptor (PRR) leucine rich repeat receptor kinases (LRR-RK) or by pattern recognition receptor-like proteins (PRR-RPs) in the plasma membrane (Wan *et al*., 2019), triggering pattern-triggered immunity (PTI). The presence of pathogen-derived effector proteins is perceived within cells by coiled-coil (CC) nucleotide binding leucine rich repeat (NLR) or Toll/interleukin 1 receptor (TIR) NLR receptors (Contreras *et al*., 2023), resulting in effector triggered immunity (ETI) (Feehan *et al*., 2020). Both scenarios share many of the same responses and indeed potentiate each other, requiring signaling through PRRs for full induction of ETI and signaling through NLRs for robust PTI (Ngou *et al*., 2021; Yuan *et al*., 2021).

Early signaling events in PTI can be studied by activating the FLAGELLIN-SENSITIVE 2 (FLS2) receptor complex, which can be elicited by the 22 amino acid epitope of bacterial flagellin (flg22), triggering several parallel protein kinase cascades, the formation of signaling lipids, such as phosphatidic acid (PA) (Kong *et al*., 2024) and the activation of calcium channels. The NADPH-oxidase, RBOHD, represents a well-studied plasma membrane target that is activated in an FLS2-dependent upon flg22 application (DeFalco & Zipfel, 2021). Elevated intracellular calcium activates calcium dependent protein kinases (CDPKs), which together with PA amplify RBOHD activation and the production of reactive oxygen species (ROS). ROS represents another quintessential signaling compound, and is an early mobile signal mediating systemic acquired resistance (SAR) (Cao *et al*., 2024). Numerous protein substrates are also phosphorylated upon FLS2-activation by mitogen-activated protein kinases (MAPKs), and the target proteins include TFs that translocate to the nucleus upon phosphorylation. In this fashion, the PAMP signal is transmitted to the nucleus, affecting gene expression. PAMP-triggered transcriptome changes can be profound, and in *Arabidopsis thaliana*, the abundance of 5189 transcripts (Hillmer *et al*., 2017) and 2689 proteins (Bassal *et al*., 2020) have been reported to respond to flg22 treatment. The TF, WRKY33, is a well-known example for a TF regulating plant immunity that is a substrate of both MAPKs and CDPKs (Liu *et al*., 2015). Upon infection of *A. thaliana* with *Botrytis cinerea*, WRKY33 directly impacts the transcription of 318 genes, thereby indirectly influencing the abundance of 2765 transcripts (Liu *et al*., 2015).

Phytohormones are among the most potent signal transducers in plants and their cross-talk shapes all aspects of plant life, including defense (Burger & Chory, 2019). A canonical defense hormone is salicylic acid (SA). Activation of the SA-receptor, NONEXPRESSOR OF PR GENES 1 (NPR1), results in NPR1 translocation to the nucleus, where NPR1 can function as a TF targeting numerous SA-dependent genes, including genes involved in SA biosynthesis. SA formation and nuclear translocation of NPR1 are both enhanced in PTI (Denoux *et al*., 2008), and 2554 genes are directly targeted (Nomoto *et al*., 2021) and 2742 differentially expressed dependent on NPR1 (Jin *et al*., 2018). Another important defense hormone is jasmonic acid (JA), and responses to JA can act both synergistically and antagonistically to SA (Hickman *et al*., 2019). JA-responses are mediated through the TF, MYC2, and its homologs, and 6175 target genes have been reported (Zander *et al*., 2020). Thus, transcriptional reprogramming by phytohormone signaling largely shapes the defense proteome.

The nexus of all signal integration is the nucleus, and the appropriate response output, i.e. the broad reprogramming of gene expression patterns, is determined by de novo synthesis as well as the translocation of TFs and other regulatory proteins into the nucleus. Spatial proteomics, the spatial resolution of protein abundance and post-translational modification on a sub-cellular scale has become technically feasible in the last few years (Mund *et al*., 2022), enabling the analysis of subcellular proteome dynamics in plants at different stages of development or upon environmental challenge, such as PAMP-treatment. One promising application is temporal and organelle-specific proximity labelling of protein populations using the TurboID system (Branon *et al*., 2018). For this technique, a genetically modified, promiscuous biotin ligase from *E. coli* is targeted exclusively to an organelle, e.g., the nucleus, where, upon exogenous application of biotin, it will biotinylate all proteins in its vicinity. The biotinylated proteins can then be identified by mass spectrometry, providing information about the subcellular distribution, e.g., in the nucleus at the time of analysis. The TurboID technology side-steps invariable pitfalls associated with traditional organelle isolation procedures, such as low yields, co-purification of other organelles or contaminants and protein leakage. Here we used a TurbeID-based approach in combination with PAMP- and control-treatments to shed light on dynamic changes in the nuclear abundance of protein regulators of gene expression during basal immunity in *Arabidopsis*.

## Materials and Methods

### Plant material

*Arabidopsis thaliana* ecotype Columbia seeds stably transformed with the R4pGWB601_UBQ10p-TurboID-YFP-NLS vector (Addgene plasmid # 127368; Supplementary figure 1) were a gift from Dominque Bergmann. The construct, cloning procedures and *Arabidopsis* lines were used previously to study the *Arabidopsis* nuclear proteome and are described in detail in (Mair *et al*., 2019).

### Plant growth

*Arabidopsis thaliana* ecotype Columbia seeds stably transformed with the R4pGWB601_UBQ10p-TurboID-YFP-NLS vector (Addgene plasmid # 127368; Supplementary figure 2) were sterilized, cold-stratified, sowed on soil and left to grow in a phytochamber under short day conditions (8 h light, 130 UML light intensity, 22°C; 16 h dark, 0 UML light intensity, 20°C) and 60 %humidity. After 2 weeks the young plants were transferred individually to separate pots. The plants were left to grow under the same conditions for another 6-7 weeks.

### Flg22, CHX and biotin treatment

Plant leaves were vacuum infiltrated with water or 2 μM flg22 or 100 µM cycloheximide (CHX) or with both flg22 and CHX and plants were left to stand for 1 h in the phytochamber. Three plants were independently treated with each treatment (water, flg22, CHX, flg22 + CHX) meaning experiments for each type of treatment were carried out in biological triplicates Then the rosette leaves (entire intact rosette) were detached from the roots quickly with a sharp cutter and rinsed with water. Rosette leaves were placed in a 1 L beaker and covered with 250 mL of 50 μM biotin solution and kept submerged for 4 h at 22°C. Afterwards, each rosette was rinsed with 250 mL of ice-cold water followed by washing with 1 liter ice cold water 3 times each for 6 min. Finally, each rosette was left shortly to dry and flash-frozen in liquid nitrogen and stored at −80°C in the late afternoon.

### Microscopy

For fluorescence microscopy, rosette leaves were cut into small pieces and placed in reaction tubes filled with water. Vacuum was applied with a syringe to remove air from the tissue. The lower part of the leaf was examined with an Axioplan2 imaging fluorescence microscope using the YFP filter. The rosette leaves were cut into discs for confocal laser scanning microscopy using an LSM 900 microscope equipped with a laser set to 488 nm with optical selection at 1.1 μM and to 410 – 545 nm for YFP detection and 635 – 640 nm for chlorophyll detection.

### Protein extraction

Rosette leaves were cryo-ground and 400 mg of ground leaf tissue were resuspended in 1.2 ml of extraction buffer (50 mM tris base, 150 mM sodium chloride, 1 % (w/v) SDS, 0.5 % (w/v) sodium deoxycholate, 1mM EGTA, 1 mM DTT, 1.5 % (v/v) protease inhibitor cocktail, pH 7.5). Homogenates were vortexed and then mixed for 10 min at 95°C then for 20 min at 22°C. 0.6 μl of lysonase was added and then the suspension was mixed further for 15 min at 22°C. Afterwards, it was sonicated for 5 min in cold water and then centrifuged at 16,000 g for 10 min at 10°C. The supernatant was transferred to a new reaction tube and centrifuged at 20,000 g for 30 min at 10°C. The supernatant was transferred to a new reaction tube and used immediately.

### SDS-PAGE and western blot

Protein extracts were separated according to their molecular weight by SDS-PAGE using 10% polyacrylamide gels. Apparent molecular weight was estimated with the blue eye prestained protein marker PS-104 (Jena biosciences) and the page ruler plus prestained protein ladder 26619 (Thermo Fisher Scientific). Proteins were blotted onto a nitrocellulose membrane using a current of 0.8 mA/cm^2^ for 1 h. The membrane was blocked with either 5% milk or 3% BSA in TBST (10 mM tris, 150 mM NaCl, 0.1% Tween 20, pH 7.5) for 1 h. Afterwards, the membrane was incubated overnight at 4°C with either 3% milk in TBST containing the Anti-GFP polyclonal antibody or 1% BSA in TBST containing the Streptavidin-POD conjugate. The membrane was washed 6 times with TBST each time for 10 min (GFP) or 6 times with TBST each time for 5 min followed by 4 times with TBS (10 mM tris, 150 mM NaCl, pH 7.5) each time for 5 min (Streptavidin). For GFP, anti-rabbit secondary antibody was added and incubated for 1 h shaking at RT. The membrane was washed as before. ECL prime western blotting detection reagent (Sigma-Aldrich) and ECL plus western blotting substrate (Thermo Fisher Scientific) were used for visualization of protein bands. Finally, the membrane was fixed on a metal cassette and a film developed using an Optimax 2010 film processor (Protec).

### Removal of excess free biotin

PD-10 gel filtration columns (Sigma Aldrich) were equilibrated five times each with 5 mL cold extraction buffer without protease inhibitor by gravity flow. 2.5 mL protein extract was applied and allowed to enter the column bed completely. Proteins were eluted with 3.5 mL of cold extraction buffer without protease inhibitor.

### Enrichment of biotinylated proteins

Dynabeads Myone Streptavidin C1 (Thermo Fisher Scientific) were washed according to manufacturer’s instructions. The beads were vortexed for 1 min and then the desired volume was transferred to a new 2 mL reaction tube. Equal volume or at least 1 mL of extraction buffer without protease inhibitor was added to the beads and mixed. The tube was placed on a magnet for 1 min to precipitate beads and the supernatant was discarded. The beads were suspended again in a volume of extraction buffer equal to the initial volume of beads. The tube was placed on a magnet for 1 min to precipitate beads and the supernatant was discarded. This was repeated twice. 16 mg of protein extract (supplemented with 1.5% (v/v) protease inhibitor) was split and applied to four 5 mL Lo-bind reaction tubes (Eppendorf) each containing 100 μL of washed beads and incubated on a rotor wheel at 4°C for 16 h. Beads were precipitated on a magnetic rack and washed subsequently as follows: (1) two times with cold extraction buffer (2) one time with cold 1 M potassium chloride (3) one time with cold 100 mM sodium carbonate (4) one time with 2 M urea in 10 mM tris pH 8 at RT (5) two times with cold extraction buffer. All washes were done with 1 mL and for 8min rotating. The beads were subsequently used without storage.

### On-bead protein digestion

1 ml of 50 mM tris pH7.5 was added to the beads and mixed on a rotor wheel for 8 min twice. The beads were transferred to a new 1.5 ml reaction tube and mixed with 1 ml 2 M urea in 50 mM tris pH 7.5 for 8 min. Beads were precipitated on a magnetic rack. 80 μL of trypsin buffer (50 mM tris pH 7.5, 1 M urea, 1mM DTT) and 2 μL of trypsin (0.2 μg/μl) were added to the beads and incubated for 3 h shaking at 800 rpm at 25°C. The supernatant was transferred to a new reaction tube and the beads were washed twice with 60 μl of trypsin buffer without trypsin. Supernatants were collected and pooled with the initial supernatant for a final volume of 200 μL. The solution was reduced in 4 mM DTT, mixing at 450 rpm at 25°C. Then the solution was alkylated in 10 mM iodoacetamide, mixing at 450 rpm in the dark at 25°C. Finally, 2.5 μL of trypsin (0.2 μg/μl) were added and the solution was incubated at 25°C shaking at 800 rpm overnight. Peptides were dried in a vacuum concentrator.

### Peptide desalting

C-18 STAGE tips were produced in-house and conditioned with 100 μl 80% ACN, 0.1% FA by centrifugation for 2 min at 1,500 g at RT. Tips were then equilibrated twice with 100 μl 0.1% FA by centrifugation for 2 min at 1,500 g at RT. The dried peptides were dissolved in 200 μl 0.1% FA, applied to the equilibrated tips and then centrifuged two times for 2 min at 1,500 g at RT. Tips were washed twice with 100 μL 0.1 % FA and centrifuged as above. Peptides eluted with 50 μl 80% ACN, 0.1% FA by centrifugation for 1 min at 1,500 g at RT in new reaction tubes. Elution was repeated and the combined eluate was dried in a vacuum concentrator.

### Liquid chromatography and mass spectrometry (LC-MS)

Dried peptides were dissolved in 12 μL of 5% ACN, 0.1% TFA and measured on a Q Exactive Plus mass spectrometer on-line with an EASY nanoLC-1000 liquid chromatography system (both from Thermo Fisher Scientific). A flow rate of 250 nl/min was used. Peptides were separated using reverse phase chemistry on an ES903 easy spray column with a length of 500 mm, an inner diameter of 75 µm and a particle size of 2 µm (Thermo Fisher Scientific) and a gradient increasing from 5% to 35% ACN, 0.1% FA in 540 min. Following separation peptides were electrosprayed into the mass spectrometer using an EASY-Spray ion source (Thermo Fisher Scientific). The spray voltage was 1.9 KV and the capillary temperature was 275°C.

A Data-Dependent Acquisition (DDA) scan strategy was used, where one MS full scan preceded up to 10 MS/MS scans of product ions from the 10 most abundant precursor ions. The MS full scan parameters were set to AGC target 3E+06, resolution 70,000 and max injection time (IT) 100 ms. The MS/MS product ion scan parameters were set to AGC target 5E+04, resolution 17,500, max IT 50 ms, dynamic exclusion duration 40 s and isolation window 1.6 m/z.

### Identification of peptides and proteins

Peptides and by inference proteins were identified by matching the MS raw data with *in silico* generated peptide and product ion m/z peak lists. The TAIR10 protein database supplemented with common contaminants (14486974 residues, 35394 sequences) was searched using Andromeda in MaxQuant version 2.0.1.0. All MaxQuant settings were left as defaults with the following exceptions. The enzyme specificity was set to trypsin/p with tolerance of 2 missed cleavages.

Carbamidomethylation of cysteine was set as a static modification and oxidation of methionine and N-terminal acetylation as a variable modification. Maximum number of modifications per peptide was set to 5. PSM and protein FDR thresholds were set to 0.01. For protein quantification unique and razor peptides were used. Data normalization was done automatically by MaxQuant implementing the MaxLFQ algorithm. The normalized LFQ intensity was used as protein quantitative index (PQI) to infer protein abundance. The LFQ minimum ratio count was set to 1 and fast LFQ was set. Match between runs and second peptides were also set. Majority protein IDs of protein groups were used, in the case multiple different proteins were present in one group, leading proteins with the highest number of identified peptides were retained. Proteins that were marked as only identified by site, reverse and potential contaminants were discarded.

### Statistical data analysis

The MaxQuant protein groups output file was imported into Perseus v2.1.3.0. Biological triplicates were grouped according to treatment condition. All protein groups that did not have a valid LFQ value in measurements of at least two of three triplicates in at least one treatment group were removed. LFQ values were log_2_ transformed. Missing values were replaced by 0. All possible pairwise Student’s t-tests between treatment groups were performed with a Benjamini-Hochberg multiples testing corrected q-value threshold of α=0.05 as a measure of statistical significance.

### Further bioinformatics analysis

Gene ontology analysis of the nuclear proteome was performed using the DAVID Bioinformatics resources 6.8 (Huang et al., 2009a, Huang et al., 2009b) using default parameters. *A. thaliana* was used as background and TAIR_ID was used as identifier. Functional annotation chart was created with threshold of count: 2 and ease: 0.1.

Subcellular location of the curated nuclear proteome was checked with SUBA4 (Hooper et al., 2017) using experimental locations inferred by fluorescent protein (FP) or MS/MS studies. LOCALIZER 1.0.4 (Sperschneider et al., 2017) was used to predict organelle subcellular localization by searching for targeting sequences such as NLS in protein primary structure and by predicting transit peptides.

LFQ values of all protein groups that had a statistically significant test result in at least one of all possible pair-wise t-tests were z-score transformed, hierarchically clustered and visualized as a heatmap using the Heatmapper website (https://heatmapper.ca). These were 328.

All protein groups that had a statistically significant test result between flg22 treated leaves and any other treated leaves (water, CHX or flg22 +CHX) were used as input to query the STRING database (https://string-db.org) with highest (0.900) confidence. These were 200. The resulting protein interaction network was expanded in a second-round query by tolerating the same number of 1^st^ shell interactors, which are not part of the input dataset, as connected nodes produced tolerating only query proteins in the original query (33 in number). All HCL cluster 3, 6 and 2 protein groups (163 in number) were used to query the STRING database in the same manner (also 33 connected nodes to expand).

The 328 protein groups with a statistically significant test result were used to query the DAVID gene ontology database (https://davidbioinformatics.nih.gov). 64 of the 328 proteins were annotated as UP_keyword biological process “Transcription” with a p-value of 1.1E-5. 5 of the 328 proteins were annotated as GOTERM biological process “regulation of circadian rhythm” with a p-value of 1.1E-3.

These 64 and 5 proteins were used to query the STRING database as above, producing 11 and 4 connected nodes in the first query respectively, to expand the network.

## Results

### TurboID can be targeted to the nucleus and identifies proteins with known nuclear residence

The efficacy of the Turbo-ID approach was tested first by determining the abundance of nuclear proteins in leaves of 8-week-old *Arabidopsis* plants grown under short-day conditions (8h light, 16 h darkness) in soil. The TurboID construct (Supplementary figure 1) was expressed from a 35S-promoter, and the biotin ligase was targeted to the nucleus of leaf cells by the strong nuclear localization signal (NLS) of simian virus 40 (SV40NLS) (Kalderon *et al*., 1984), as was previously used to determine nuclear proteome composition (Mair *et al*., 2019). The presence of the fusion protein in the transgenic lines used was verified by immunodetection (Figure 1A). Nuclear targeting of the expressed fusion protein was successful, as evident from the nuclear localization of the fluorescence-tagged protein in Arabidopsis leaf epidermal cells, imaged by confocal microscopy (Figure 1B). Following treatment with biotin for 4 h and enrichment of biotin labeled proteins (Figure 1C), the number of biotinylated proteins was substantially greater in transgenic lines expressing the TurboID construct, compared to non-transformed controls with only a fraction of proteins detected. LC-MS analysis in biological triplicates identified 2105 protein groups with high confidence (i. e., with at least one unique peptide and PSM, peptide and protein FDR thresholds of α = 0.01, henceforth referred to as proteins; Table 1, Supplementary file). Tissue extracts of non-transformed plants served as negative controls for non-specific binding of proteins to the streptavidin beads, and only 104 proteins were determined for these controls (Table 2, Supplementary file), indicating that physiological biotinylation was negligible (Figure 1C). Overall, 81 proteins displaying unspecific binding were subtracted as background from the set of proteins positively identified by TurboID.

**Figure 1.**
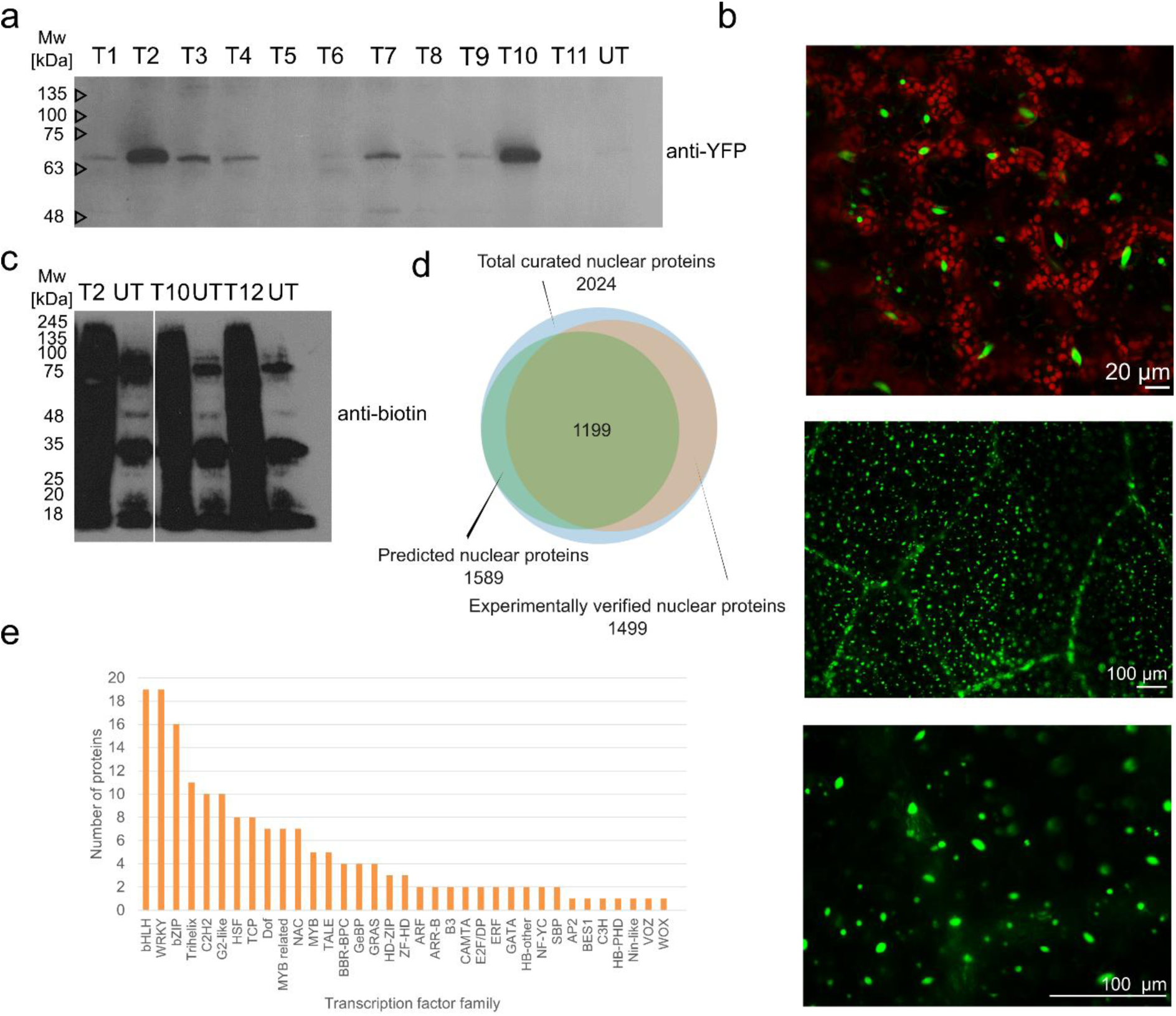
TurboID facilitated labeling and isolation of the nuclear proteome. a. Anti-YFP western blot showing the expression of the TurboID-NLS-YFP fusion protein in total leaf extract using a polyclonal anti-GFP antibody. The transgenic protein has a calculated molecular mass of around 65 kDa. Extracts from 11 transgenic plants (T1 to T11) and untransformed (UT) are shown. Figure is representative, only plants expressing the protein at levels as detected in T2 or T10 were used for subsequent experimentation. b. Nuclear localization of the TurboID-NLS-YFP fusion protein. Leaf undersides were examined under a confocal laser scanning microscope using 410-545 nm wavelengths for YFP and 635-640 nm wavelengths for chlorophyll detection. Images are representative. Scale bars, as indicated. c. Anti-biotin western blot showing protein biotinylation in total leaf extract using a Strepdavidin-Pod conjugate antibody. d. Subsets of the TurboID isolated nuclear proteome that were predicted or experimentally verified to be nuclear localized using DAVID Bioinformatics resources 6.8, SUBA4 and LOCALIZER 1.0.4. e. Exemplary classification of transcription factors identified in the nuclear proteome according to families by DAVID Bioinformatics resources 6.8.

Of the remaining 2024 proteins (Table 3, Supplementary file), 1889 proteins are predicted or have previously been determined experimentally to be located in the nucleus (Figure 1 D). Among these, 322 proteins classify as TFs or proteins related to transcription (Table 4 and Table 5, Supplementary file). These included low or transiently abundant members of the basic helic-loop-helix (bHLH), basic leucine zipper (bZIP) or WRKY families, and others such as ethylene response factors (ERF), myeloblastosis TFs (MYBs) or heat shock factors (HSFs), all known to regulate developmental and physiological processes in plants (Figure 1E). The deep and specific coverage of the nuclear proteome included detection of numerous low abundant proteins and TFs and made a quantitative study of gene expression regulators in the nucleus in PTI seem promising. Importantly, the experiments demonstrate that in our hands the TurboID-MS approach yielded a comprehensive list of bona fide nuclear proteins, enabling further analyses of dynamic changes in the nuclear proteome upon flg22 treatment.

### Nuclear proteome dynamics upon flg22 challenge

To expand our analysis to *Arabidopsis* basal immunity, rosette leaves of transgenic plants were subjected to four different treatments: (i) water, (ii) flg22, (iii) water plus the translation inhibitor cycloheximide (CHX), or (iv) flg22 plus CHX. Treatments were applied for 1 h prior to harvest and the addition of exogenous biotin and subsequent proteomic LC-MS analysis. The premise behind the experimental design was to identify and distinguish proteins that accumulate in the nucleus upon elicitation of PTI as a result of either transcriptional reprograming followed by translation and nuclear protein import (water vs. flg22 vs. flg22 plus CHX); by translocation of preexisting proteins into the nucleus (water vs. CHX vs. flg22 plus CHX); or by stabilization or degradation of proteins already present in the nucleus (also water vs. CHX vs. flg22 plus CHX). Non-transformed plants were included as negative controls, and proteins identified by LC-MS in these controls were subtracted from the TurboID set as above (Table 6, Table7 and Table 8, Supplementary file).

Upon filtering of low intensity protein signals, 1778 proteins were quantified in LC-MS measurements of biological triplicates of the differently treated rosette leaves (Table 9, Supplementary file). Pair-wise t-testing of all experimental conditions uncovered 328 proteins with a significant difference in abundance between at least two of the four experimental conditions with a Benjamini-Hochberg false discovery rate (FDR) threshold of α=0.05 (Table 10, Supplementary file).

Hierarchical clustering was used to classify the differentially abundant proteins (Figure 2), and the proteomic data suggest at least six clusters of proteins that change differentially over time with different treatments.

**Figure 2.**
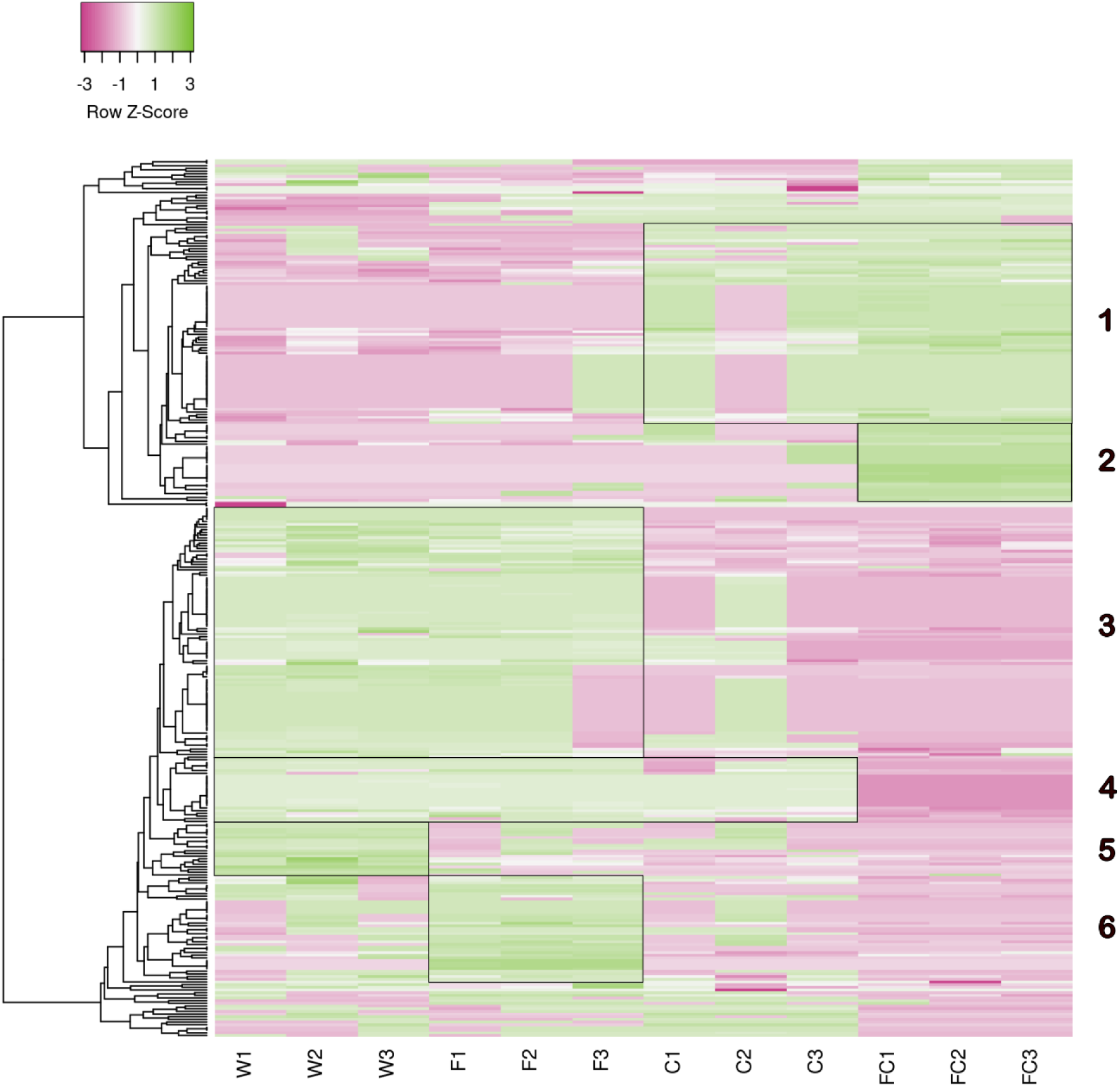
Hierarchical clustering of the abundance of all 328 nuclear proteins with a significant test result (Benjamini-Hochberg corrected p-value < 0.05) between at least two of the four tested conditions (mock treated W, flg22 treated F, CHX treated C and flg22 and CHX treated FC). Clusters are indicated and numbered.

Proteins in cluster 1 accumulated in the nucleus in response to CHX, and perhaps also flg22, conceivably as a result of perturbed turn-over and degradation of cognate repressors, or of missing protein components of the degradation or nuclear export machinery (Table 11, Supplementary file).

A clearly different pattern is displayed by proteins in cluster 2, demarking proteins that were increased in nuclear abundance upon treatment with flg22 plus CHX. These proteins may have translocated to the nucleus but were not newly synthesized after initiation of immunity, again possibly due to depletion of negative regulators by co-treatment of flg22 with CHX (Table 12, Supplementary file).

Reciprocally, proteins representing cluster 3 were decreased in nuclear abundance in the presence of CHX, regardless of whether or not flg22 was applied. Proteins in cluster 3, thus, represent flg22-independent nuclear proteins of possibly high turnover, which depleted rapidly when their de novo biosynthesis was blocked. (Table 13, Supplementary file).

Proteins in cluster 4 were similarly depleted from the nucleus when CHX was applied, but only upon co-application of flg22. This pattern suggests that proteins in cluster 4 might display an equilibrium of high turnover and resynthesis and are additionally destabilized upon flg22 challenge.

Proteins of cluster 5 are characterized by a decreased nuclear abundance upon flg22 challenge and an overall reduced nuclear abundance in CHX-treated plants as compared to water-treated plants. This pattern indicates that cluster 5 proteins are subject to destabilization and/or display nuclear export upon flg22 challenge, both of preexisting and newly synthesized protein populations.

Finally, cluster 6 comprises proteins that displayed enhanced nuclear abundance upon flg22 challenge, while showing a reduced nuclear abundance in the presence of CHX. These proteins were newly synthesized and imported into the nucleus upon exposure of leaves to flg22 and are likely subject to rapid turnover (Table 14, Supplementary file).

The comparison of the proteomic datasets by hierarchical clustering analysis (Figure 2) indicates that the TurboID-MS analysis of nuclear proteomic changes with flg22 challenge can identify the dynamic behavior of a substantial number of nuclear proteins. Moreover, the comparative analysis informs about the respective contributions of *de novo* biosynthesis, destabilization or translocation that in sum define nuclear protein homeostasis.

### Dynamic patterns of TF abundance

With around 8% of protein coding ORFs encoding TFs in eukaryotic genomes (Pruneda-Paz *et al*., 2014) and more than 2600 loci classified as TFs in *Arabidopsis* (Mitsuda & Ohme-Takagi, 2009), these proteins profoundly impact gene expression by exercising transcriptional control. Among many others, numerous bHLH, bZIP and WRKY TFs which diversify their regulation of target genes by homo- and heteromeric di-and multimerization, were differentially abundant in leaf nuclei upon flg22 treatment (Figure 3). Examples that were more abundant in the nucleus upon flg22 treatment include bHLH129 (Tian *et al*., 2015) which contributes to ABA responses potentially impacting stomatal immunity; or bHLH13/JAM2, which negatively impacts JA mediated defenses (Song *et al*., 2013). MYC2, a nuclear protein tightly controlled by proteasomal degradation (Chico *et al*., 2020), was not detected in water and flg22 treated, but accumulated in CHX treated and flg22 plus CHX co-treated nuclei. This pattern indicates relief of cognate repressor activity by way of translation and protein turn-over inhibition, underscoring control of MYC2 activation and the JA response in PTI. WRKY TFs are unique to plants and have well documented roles in both biotic and abiotic stress responses. Known constitutively expressed and PAMP induced WRKY TFs (Birkenbihl *et al*., 2018) were detected in leaf nuclei, many of the latter were more abundant after exposure to flg22 (Figure 3). These included WRKY33 and WRKY40, preeminent regulators of PTI (Saha *et al*., 2023; Chen & Zhang, 2024) as well as WRKY6 which has been implicated downstream of NPR1 in SA mediated leaf senescence and pathogen defense (Robatzek & Somssich, 2002; Zhang *et al*., 2021) and WRKY72, described to attenuate JA signaling in rice (Hou *et al*., 2019).

**Figure 3.**
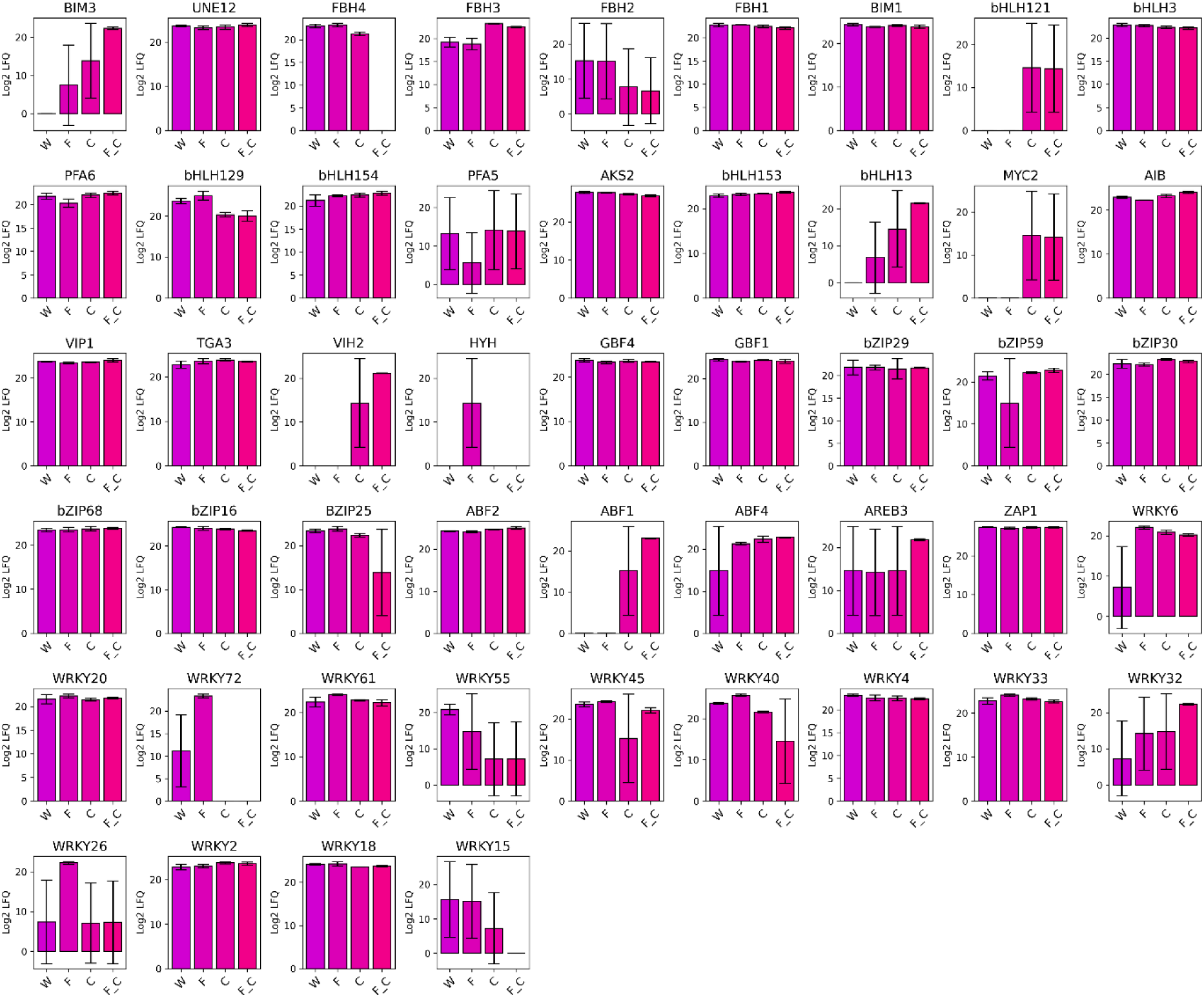
bHLH, bZIP and WRKY transcription factors in the set of all 328 nuclear proteins with a significant test result (Benjamini-Hochberg corrected p-value < 0.05) between at least two of the four tested conditions (mock treated W, flg22 treated F, CHX treated C and flg22 and CHX treated FC). Their abundance in the four experimental treatment conditions (mock treated W, flg22 treated F, CHX treated C and flg22 and CHX treated FC) is shown in the bar charts.

### Prediction of interaction networks supports hierarchical clustering

Proteins often exert their activity as components of homo- or heteromeric (multi-)functional units or complexes. To help pin-point potential regulatory modules imported into the nucleus that reprogram gene expression and may orchestrate PTI, proteins in clusters 2, 3 and 6 as well as those with a significant difference in their abundance between flg22 treatment and any of the remaining three experimental conditions were queried for interactions using the Search Tool for the Retrieval of Interacting Genes/Proteins (STRING) database (Szklarczyk *et al*., 2023). Only matches with the highest (0.900) level of confidence were included in our representation. The input sets produced overlapping, highly significant output sets of functionally diverse protein interaction networks, many more than might be attributed to chance (Figures 4 and 5; Table 15 and Table 16, Supplementary file; Supplementary figure 2). Proteins classified as related to transcription and circadian rhythms by gene ontology analysis (64 and 5, respectively) were also queried accordingly (Table 17 and Table 18, Supplementary file; Supplementary figure 2). Querying cluster 1 entries, proteins that were abundant in the nucleus as a result of cycloheximide exposure, did not produce any interactions beyond what would be expected from a random input set.

**Figure 4.**
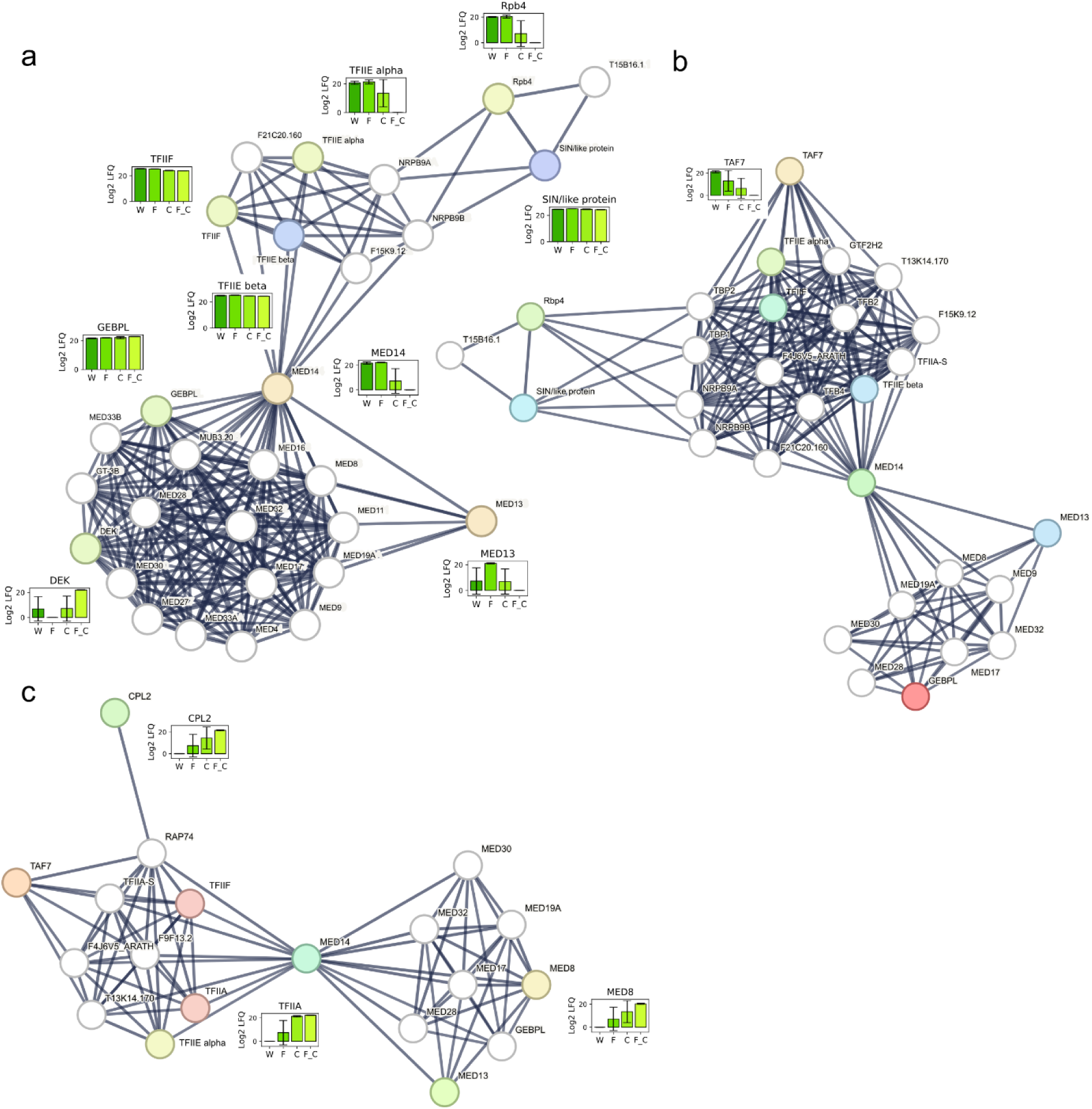
Protein interaction networks of mediator complex components produced with the STRING database with highest (0.900) confidence. Colored nodes represent proteins that were measured in the leaf nuclei so were part of the input set, their abundance in the four experimental treatment conditions (mock treated W, flg22 treated F, CHX treated C and flg22 plus CHX treated FC) is shown in the bar charts. White nodes were added by STRING to expand the network in a second query that tolerated an equal number of secondary interactors as primary interactors produced by the original query of the input set. a. input set was all proteins in clusters 2, 3 and 6. b. input set was all proteins with a significant difference in their abundance between flg22 treatment and any of the remaining three experimental conditions. c. input set was all proteins characterized as related to transcription by DAVID Bioinformatics resources 6.8.

**Figure 5.**
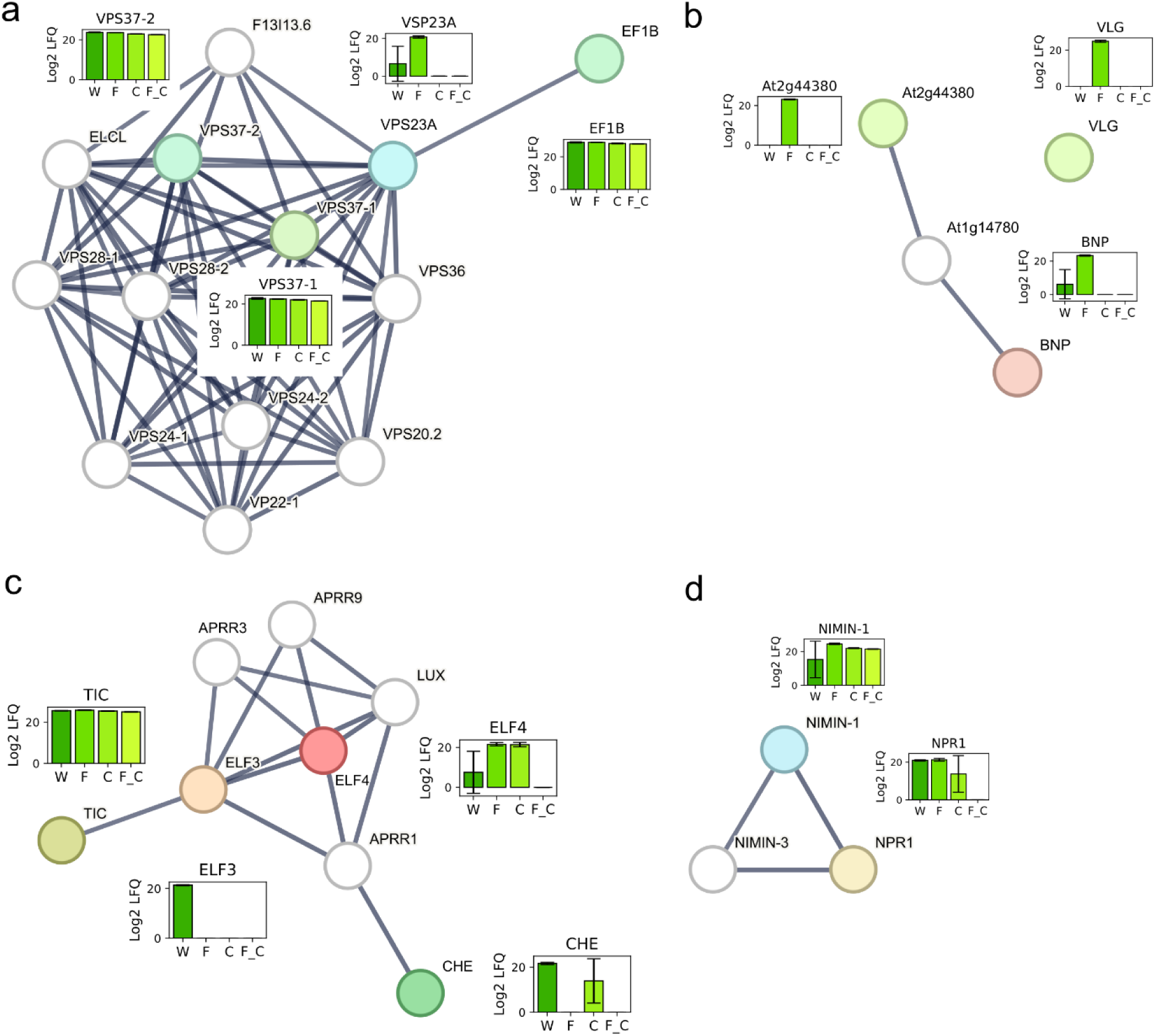
Protein interaction networks of a. ESCRT complex components, b. cysteine/histidine rich proteins, c. circadian clock components, d. NPR1 and associated proteins produced with the STRING database with highest (0.900) confidence. Colored nodes represent proteins that were measured in the leaf nuclei so were part of the input set, their abundance in the four experimental treatment conditions (mock treated W, flg22 treated F, CHX treated C and flg22 plus CHX treated FC) is shown in the bar charts. White nodes were added by STRING to expand the network in a second query that tolerated an equal number of secondary interactors as primary interactors produced by the original query of the input set. a. input set was all proteins in clusters 2, 3 and 6. b. all proteins in clusters 2, 3 and 6 and all proteins with a significant difference in their abundance between flg22 treatment and any of the remaining three experimental conditions as input sets produced this network. c. input set was all proteins characterized as related to circadian rhythms by DAVID Bioinformatics resources 6.8. d. input set was all proteins with a significant difference in their abundance between flg22 treatment and any of the remaining three experimental conditions.

### Mediator complex

For instance, the mediator complex contains 29 conserved subunits in *Arabidopsis* organized into head-, middle- and tail modules. The mediator complex can be considered a part of the basal transcription machinery because it is essential for all aspects of RNA polymerase II (Pol II) mediated transcription (Freytes *et al*., 2024) and coordinates the assembly of the preinitiation complex (PIC) together with the general TFs TFIID and TFIIH, recruiting Pol II and TFIIA, TFIIB, TFIIE and TFIIF. Importantly, the mediator complex modulates TF activity and impacts on the expression of TF-specific gene sets. Concurrently, in our STRING analysis, TFIID, TFIIE and TFIIF as well as Pol II and Pol III subunits were associated with the mediator complex and were found more abundant in nuclei of untreated and PTI elicited leaves compared to leaves experiencing translational inhibition (Figure 4). Mediator 8 is a head module subunit which positively affects resistance to *A. brassicicola* (Kidd *et al*., 2009). Mediator 14 forms a backbone necessary for complex assembly, and several mutant alleles are not viable when homozygous (Zhang *et al*., 2013). Mediator 14 has a documented function in SA-mediated immunity and in SAR (Zhang *et al*., 2013) and was found in our analysis to be abundant in the nucleus of untreated and flg22 stimulated leaves (Figure 4a). Depletion of the mediator 14 subunit was observed upon CHX exposure, suggesting the protein is turned over rapidly and was not replenished (Figure 4a). Mediator 13 is part of a separable kinase module (CKM) that regulates mediator function both positively and negatively by competing for binding with Pol II and can phosphorylate various members of the PIC. Mediator 13 was significantly most abundant upon flg22 challenge alone (Figure 4a), indicating expression and nuclear import in PTI. Increased mediator 13 dependent expression of the SA marker gene PR1 and resistance to necrotrophic and bacterial pathogens have previously been reported (Zhu *et al*., 2014).

### ESCRT I complex subunits moonlighting in the nucleus

Endosomal sorting complex required for transport (ESCRT) is a dynamically assembled heteromeric multi-subunit protein complex essential for trafficking membrane spanning as well as cytosolic ubiquitinated cargo by way of multi vesicular bodies (MVBs) to the vacuole for degradation (Gao *et al*., 2017). ESCRT-I component VPS37-1 plays a role in internalization of the flg22 receptor FLS2 and stomatal immunity (Spallek *et al*., 2013) which is also linked to ABA signaling. Furthermore, ESCRT-I components VPS23A and FREE1 dependent endosomal sorting of cytoplasmic ubiquitinated ABA receptors PYR1/PYL-RCAR effects their turn-over and the ABA response.

ESCRT subunits showed conspicuous changes in nuclear abundance upon treatment with the elicitors. ESCRT subunits are primarily membrane associated, yet the ESCRT-I subunits VPS37-1 and VPS37-2 as well as VPS23A were abundant in leaf nuclei, the former two constitutively under all treatments, the latter explicitly after exposure of leaves to flg22 (Figure 5a). Interestingly, an alternate function of the protein FYVE-Domain Protein Required for Endosomal Sorting 1 (FREE1) in the nucleus has previously been described, and FREE1 directly inhibits the DNA-binding activity of ABA responsive transcriptional activators ABF4 and ABI5, thereby attenuating ABA signaling (Li *et al*., 2019). The results suggest that VPS23A may be moonlighting in the nucleus with a particular function in PTI.

### Do cysteine/histidine rich proteins regulate basal immunity in the nucleus?

The TurboID-MS analysis of nuclear proteome changes in flg22 treated *Arabidopsis* leaves revealed a number of currently little understood proteins displaying pronounced increases in their nuclear abundance. Examples include the proteins VACUOLE-LESS GAMETOPHYTE (VLG), BINUCLEATE POLLEN (BNP) and the non-annotated gene product of locus At2g44380, which all accumulated exclusively in nuclei of flg22 challenged leaves, indicating *de novo* translation and nuclear import upon PTI elicitation (Figure 5b).

All three proteins are members of a protein family containing divergent C1 (DC1) domains. DC1-domain containing proteins are only found in plants and comprise a family of 140 members whose function remains currently obscure (D’Ippolito *et al*., 2017). As cysteine/histidine rich domains, DC1-domains feature a conserved primary structure signature that coordinates two zinc ions and may mediate protein/protein interactions, protein membrane tethering and DNA and RNA binding. VLG (At2g17740) (D’Ippolito *et al*., 2017) and BNP (At2g44370) (Arias *et al*., 2023) were both shown to be essential for male and female gametogenesis, and homozygous mutant progeny for both genes was not viable. The genes are part of a family clade that also contains At2g44380, which has been implicated in the salt stress response (Gao *et al*., 2016), and are homologous to *Capsicum annum* (pepper) CaDC1, whose silencing resulted in elevated susceptibility to biotrophic pathogens and reduced SA accumulation up to 36 hours post infection with virulent strains in pepper and *Arabidopsis* (Hwang et al., 2014).

At the subcellular level, VLG and BNP were previously detected in the late endosome and in multi-vesicular bodies (MVBs) in the young sporophyte and developing gametophyte, but not in the nucleus. BNP and At2g44380 potentially interact with one of four MACPF domain containing proteins in *Arabidopsis*, At1g14780 (Gao *et al*., 2016) (Figure 5b). This interaction may contribute to the regulation of SA mediated defenses based on the similarity of the At1g14780-gene product with CAD1, another MACPF domain containing protein (Morita-Yamamuro *et al*., 2005).

In sum, based on their flg22-dependent nuclear abundance, the detected DC1 domain containing proteins may exert an as of yet undiscovered role in immunity to biotrophic pathogens in the nucleus of the mature *Arabidopsis thaliana* sporophyte. As in pepper, DC1 domain liaised nucleic acid binding may be quintessential to triggering defense signaling downstream of PAMP perception in the nucleus. Alternatively, DC1-proteins may facilitate scaffolding and nuclear translocation of immunity regulators, such as MACPF domain containing proteins, suggesting potential moonlighting function in addition to their endosomal/MVB roles in gametophyte development and the young sporophyte.

### Circadian clock and evening complex proteins in PTI

The circadian clock, a genetic oscillator composed of several interconnected transcriptional-translational feedback loops (TTFL), attunes many aspects of plant physiology to a 24-hour time period (Lu *et al*., 2017). The circadian clock is influenced in part by environmental input cues or *zeitgebers,* and output pathways are gated in a time dependent manner. The core TTFLs are the morning loop composed of the proteins, CIRCADIAN CLOCK ASSOCIATED 1 (CCA1) and LATE ELONGATED HYPOCOTYL (LHY) and the evening loop, that encodes the evening complex (EC), composed of EARLY FLOWERING 3 (ELF3), ELF4 and LUX ARRHYTHMO (LUX) (Huang & Nusinow, 2016). As physiological control is exercised primarily by regulation of transcription, many clock components are TFs and display nuclear localization.

Four clock components, TIME FOR COFFEE (TIC), ELF3, ELF4 and CHE were all detected by TurboID-MS in leaf nuclei of untreated plants (Figure 5c). TIC is constitutively localized in the nucleus irrespective of time of day, functioning close to the core TTFLs to maintain rhythmicity and amplitude of the expression of their components, particularly *LHY*, *ELF3* and *ELF4* (Ding *et al*., 2007). In line with these previous findings, in our analysis TIC was equally abundant in nuclei of both flg22 treated and CHX treated leaves and did not display indications of protein turnover.

EC proteins accumulate in the late-afternoon and evening (Lu *et al*., 2017), and we observed ELF3 and ELF4 in nuclei of untreated leaves (Figure 5c). Knockout mutations in all EC genes lead to arrythmia of the clock itself (Huang & Nusinow, 2016) and mutations in *ELF3* have been shown to compromise stomatal immunity and ROS production (Lai *et al*., 2012). In our TurboID-MS analysis, ELF3 was depleted from nuclei upon flg22 challenge and CHX treatment, suggesting ELF3 turn-over and impaired assembly and function of EC in PTI. This assumption is based on the notion that ELF3 is recruited to the nucleus by ELF4 (Herrero *et al*., 2012), is necessary for EC assembly (Nusinow *et al*., 2011), and that its function is dependent on its nuclear localization (Herrero & Davis, 2012; Anwer *et al*., 2014). ELF4 increased in abundance in nuclei of PTI-induced leaves (Figure 5c), possibly stabilizing EC binding to DNA (Silva *et al*., 2020).

### Regulators of the SA response in PTI

The circadian clock is integrally linked to the key signaling pathways of plant immunity. The TF CHE, next to its role in the clock as a central oscillator component regulating the morning loop by way of CCA1 expression (Pruneda-Paz *et al*., 2009), binds the promoter region in the *isochorismate synthase 1 (ICS1)* gene involved in SA biosynthesis, thereby determining the circadian oscillation of the *ICS1* transcript and SA levels (Zheng *et al*., 2015). At the protein level, the CHE TF was approximately equally abundant in leaf nuclei of mock and CHX treated plants, yet conspicuously absent when leaves were treated with flg22, both in the presence or absence of CHX (Figure 5c). This pattern suggests the CHE protein was specifically depleted from the nucleus upon flg22 treatment, either by way of protein export or by degradation.

In previous SAR experiments, CHE was directly responsible for the induction of *ICS1* transcription, SA accumulation, defense gene expression and the onset and establishment of SAR in distal plant tissue. However, CHE did not play a role in immunity at the infection site, when plants were locally infiltrated with avirulent or virulent pathogens (Zheng *et al*., 2015; Cao *et al*., 2024). Here, exposure of the entire leaf to flg22 strictly induced PTI throughout the tissue, a scenario where full-scale induction of SA mediated defenses as in SAR may be detrimental. Hence, depletion of CHE from the nucleus may represent an additional safeguard from SAR induction in local tissue (Cao *et al*., 2024).

The transcriptional co-activator NON-EXPRESSOR OF PATHOGENESIS-RELATED GENES 1 (NPR1, also known as NON-INDUCIBLE IMMUNITY 1 (NIM1)) is one of three SA receptors and a central protein coordinating all layers of immunity to biotrophic pathogens (Fu & Dong, 2013; Zavaliev & Dong, 2024). NPR1 is required for full induction of flg22-induced basal immunity, avirulent pathogen induced ETI and is indispensable for broad-spectrum systemic resistance (Liu *et al*., 2020), the *npr1* mutant being completely compromised in SAR (Cao *et al*., 1994). The NPR1 protein is also intrinsically connected to the circadian clock, fortifying it to gate defense towards the morning under SA induced perturbation of cellular redox rhythm (Zhou *et al*., 2015).

Subcellular localization and protein turn-over are inherent to NPR1 activity and function (Mou *et al*., 2003; Spoel *et al*., 2009). In our TurboID-MS experiments, NPR1 was detected in the nuclei of mock treated but less so in nuclei of CHX treated leaves (Figure 5d), in agreement with constitutive nuclear import and degradation of the monomer to suppress activation of defense gene expression and SAR (Spoel *et al*., 2009). The flg22 treatment led to a slight increase in nuclear NPR1 protein abundance but not to alleviation of turn-over or stabilization, as evident from a lack of NPR1 in nuclei of the leaves treated with both, flg22 and CHX. Similar, classic experiments wherein SA treatment and avirulent pathogen challenge increased *WRKY* and *PR-1* gene expression and induced SAR yet also did not abolish NPR1 degradation in the nucleus are reminiscent (Spoel *et al*., 2009).

The functionally related protein, NIM1-INTERACTING 1 (NIMIN-1), was also most abundant in nuclei of leaves treated with flg22 and was also present in nuclei of all other treated leaves (Figure 5d), suggesting elevated translation and nuclear import upon flg22-challenge. NIMIN1 is one of four related NIM1 interacting proteins that all modulate the expression of PATHOGENESIS-RELATED 1 (PR-1), which is a key mediator of SAR (Hermann *et al*., 2013). NIMIN-1 has been shown to interact with NPR1/NIM1 and is a major negative regulator of SAR and ETI, over-expressor lines being unable to mount the former and severely compromised in the latter, similar or more so respectively, than the *npr1* mutant (Weigel *et al*., 2005).

## Discussion

The TurboID technique has sparked phytologist’s interest in recent years (Xu *et al*., 2023), because of its ability to identify protein-protein interactions and proximal interactomes of specific targets *in vivo*. Transgenic bait proteins generally retain activity, and the biotin-labeling reaction is bio-compatible and proceeds in the scale of minutes. Biotinylated prey can be isolated and purified under stringent conditions, ideal for down-stream mass spectrometry (MS). Several recent applications of TurboID approaches include mapping plant immune signaling pathways (Zhang *et al*., 2019; Kim *et al*., 2023; Powers *et al*., 2024), the meiotic chromosome axis (Feng *et al*., 2023) and applications coupled to Split-YFP (Huang *et al*., 2025) and CRISPR (Zhang *et al*., 2026). On the down-side, *in planta* many transient interactions may only last for seconds, the distance spanned by TurboID labeling is between 100 and 350 Å (Mair & Bergmann, 2022), a ballpark in the structural biology world, and the identification of proteins by subsequent MS-analysis is of course indirect. Chemical or metabolic (photo-) cross-linking, where reactions proceed within seconds, Cα distances are typically between 0 and 35 Å, and MS identification of two cross-linked peptides from interacting proteins is direct (Piersimoni *et al*., 2022) could be an advantageous approaches to TurboID.

While these issues must be considered for the interpretation of any TurboID experiment, they are not as prevalent when TurboID is used to exclusively label and isolate sub-cellular proteomes as done here and previously (Mair *et al*., 2019). Specific interactions of cellular proteins proximal to the transgenic target were not the focus, instead an NLS directed the biotin ligase to the nucleus, where it labeled essentially all proteins enclosed by the nuclear envelope irrespective of their spatial relationship to one-another. Efficient purification yielded a pure and abundant nuclear protein fraction amenable to conventional discovery proteomics approaches from individual plants. More than 90% of the detected >2000 proteins were *bona fide* nuclear proteins, attesting to the power of TurboID in this context. Common confounders associated with conventional biochemical or biophysical nuclear enrichment procedures might have impacted the interpretation of previous studies of the nuclear proteome in immunity in different plant species, which in part employed highly artificial systems (Howden *et al*., 2017; Narula *et al*., 2019; Rajamaki *et al*., 2020; Ayash *et al*., 2021). By contrast, the comparatively large amounts of required biological starting material, co-isolation of other organelles and impurities, or damaged or ruptured nuclei and protein leakage were all easily avoided by applying the TurboID approach.

The alleviation of transcriptional repression by way of repressor protein degradation is a common theme in the control of gene expression, as exemplified for instance by the perception of phytohormones, such as ABA, gibberellic acid (GA), JA, auxin and also SA, by their cognate receptors. Translation inhibition with CHX was used to study flg22-induced changes in protein turn-over in the nucleus and translation initiation followed by protein shuttling to the nucleus. Absence in nuclei of CHX treated leaves indicated rapid protein turn-over. Protein abundance in nuclei of PAMP treated together with absence in nuclei of co-treated leaves indicated protein shuttling upon induction of basal immunity or constant turn-over and clearance of the protein in the nucleus, as was the case for NPR1 (Figure 5c). Previous research has shown, that the transcription of the vast majority of genes induced by flg22 was also induced when plants were co-treated with CHX and even with CHX alone, highlighting the key role of protein repressors and their turn-over in the plant defense response (Navarro *et al*., 2004).

In nature plants are exposed to a variety of pathogens with different life styles, and the overlapping pathways that constitute robust immunity, PTI, ETI and SAR, will all concomitantly be active and interconnect in some form. Exposing entire plants to high concentrations of flg22 to elicit strong PTI was clearly a somewhat unnatural scenario. Nevertheless, the chosen experimental design enabled the distinction of nuclear protein import and turn-over in basal immunity strictly downstream of FLS2.

Several proteins with documented functions in endosomal sorting and localized in endosomes and MVBs, VPS23A, and the cysteine/histidine rich proteins BNP, VLG and the At2g44380 gene product, were synthesized and imported into the nucleus after stimulation by the PAMP, suggesting a regulatory function in basal immunity. SAR signaling is considered to be initiated following induction of ETI, so downstream of intracellular NLR receptors, which were not triggered. One may speculate on possible control mechanisms of SAR induction which may be deleterious to plant fitness in the context of basal immunity alone, such as the depletion of CHE or the increased import of NIMIN-1 in response to flg22.

To conclude, remodeling of the nuclear proteome as a result of protein import, degradation and changes in turn-over was characterized comprehensively, capturing substantial numbers of low or transiently abundant regulators of gene expression. Undoubtedly the data can be mined further to uncover other relevant facts in addition to those reported here, which can be verified and expanded on with subsequent genetic and biochemical experiments.

## Acknowledgments

M.A. was funded by the Deutsche Forschungsgemeinschaft (DFG) Research Training Group RTG 2498 “Communication and Dynamics of Plant Cell Compartments”, grant number 400681449/GRK2498. W.H. was supported by the Deutsche Forschungsgemeinschaft (DFG – 514901783) through the Collaborative Research Centre 1664 “Plant Proteoform Diversity-SNP2Prot”, project B02, as well as by the Leibniz Association. All other authors thank the Martin-Luther University Halle-Wittenberg and the Leibniz Association for core funding. We thank Dennis Psaroudakis for depositing the data at ProteomeXchange/PRIDE as well as with help in ARC construction.

## Author contributions

M.A., J.L, I.H. and W.H. designed the research. M.A. performed the experiments, C.P. performed the LC-MS analysis, D.T. assisted with plant growth and biochemistry experiments, N.B. assisted with the biochemistry experiments and microscopy, M.A. and W.H analyzed the data. W.H. and I.H. wrote the paper.

## Data availability

The mass spectrometry proteomics data have been deposited to the ProteomeXchange Consortium via the PRIDE (Perez-Riverol *et al*., 2025) partner repository with the dataset identifier PXD078376

---

**Supplementary figure 1.**
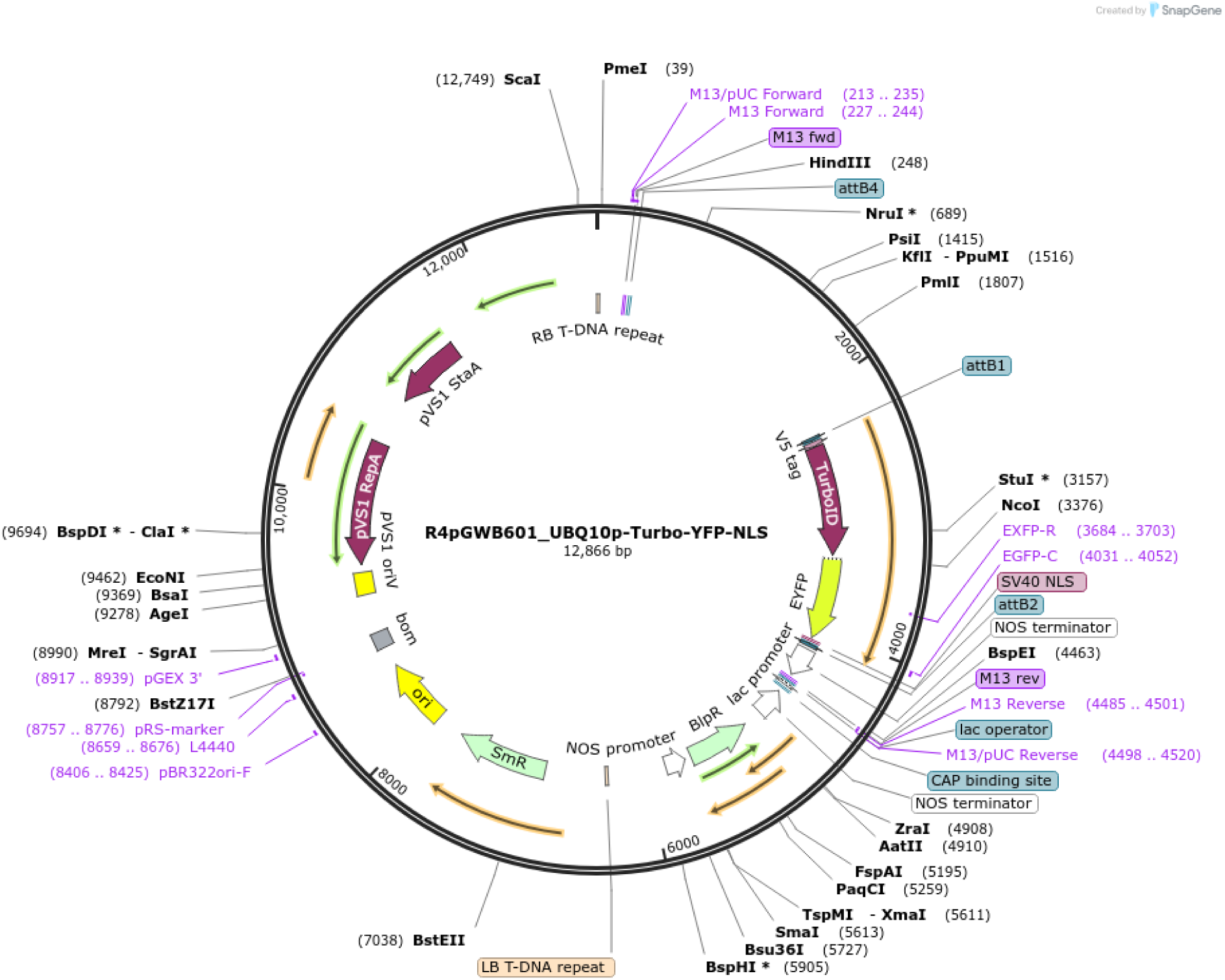
Full Sequence Map for R4pGWB601_UBQ10p-Turbo-YFP-NLS

**Supplementary figure 2.**
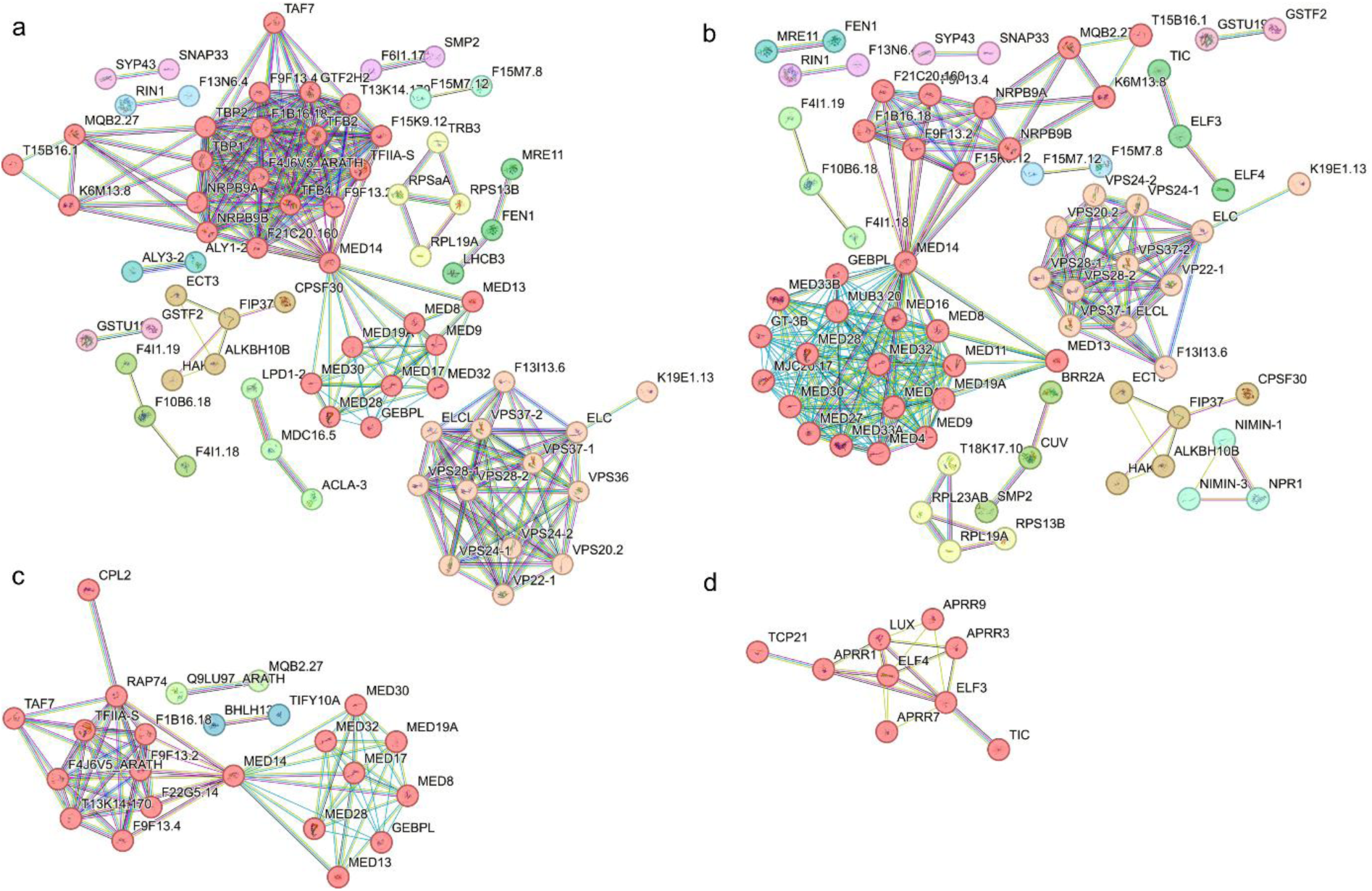
K-means clustering and functional annotation of protein interaction networks produced by the STRING database for input sets a. all proteins in clusters 2, 3 and 6. b. all proteins with a significant difference in their abundance between flg22 treatment and any of the remaining three experimental conditions. c. all proteins characterized as related to transcription by DAVID Bioinformatics resources 6.8. d. all proteins characterized as related to circadian rhythms by DAVID Bioinformatics resources 6.8. Cluster descriptions are found in Supplementary file 1, Tables 15 to 18.

